# Analysis of female song provides insight into the evolution of sex differences in a widely studied songbird

**DOI:** 10.1101/2020.03.28.013433

**Authors:** Matthew R. Wilkins, Karan J. Odom, Lauryn Benedict, Rebecca J. Safran

## Abstract

Understanding the patterns and processes related to sexual dimorphism and sex differences in diverse animal taxa is a foundational research topic in ecology and evolution. Within the realm of animal communication, studies have traditionally focused on male signals, assuming that female choice and male-male competition have promoted sex differences via elaboration of male traits, but selection on females also has the potential to drive sexual differentiation in signals. Here, we describe female song in barn swallows (*Hirundo rustica erythrogaster*) for the first time, report rates of female song production, and couple song data with plumage data to explore the relative degree to which sex differences in phenotypic traits are consistent with contemporary selection on males versus females. During previous intensive study of male song over two years, we opportunistically recorded songs for 15 females, with matched phenotypic and fitness data. We randomly selected 15 high-quality samples from our larger male dataset to test whether sex differences in song and plumage are more strongly associated with fledgling success for females or genetic paternity for males. Analyses included 35 potential sexual signals including 22 song parameters and 13 plumage traits. Outcomes indicate that: female songs were used in multiple contexts, restricted primarily to the beginning of the breeding season; song traits showed greater sexual differentiation than visual plumage traits; and trait correlations with reproductive success in females, rather than males, predicted sex-based differences in song and plumage. These results are consistent with phylogenetic studies showing that sex-based phenotypic differences are driven by changes in females, highlighting the potential role of female trait evolution in explaining patterns of sexual differentiation. To achieve a better understanding of sex differences and dimorphism, we require comprehensive studies that measure the same traits in males and females and their fitness consequences.

## INTRODUCTION

Ecologists and evolutionary biologists have long sought to understand the processes driving dimorphism and other sex-based phenotypic differences (Andersson, 1994; Badyaev & Hill, 2003; Burns, 1998; Darwin, 1859, p. 94; Darwin, 1801, p. 396; Hedrick & Temeles, 1989; Lande, 1980; Ng et al., 2019). However, owing to historical biases, studies of the drivers of differentiation in sexual signaling traits have traditionally focused on male signals, and most approaches assume that sexual selection has promoted sex differences via elaboration of male traits (Badyaev & Hill, 2003; Freed, 2000; Langmore, 1998; Riebel, 2016; Riebel et al., 2005, 2019; Rosvall, 2011). Yet, meta-analyses of the strength of sexual selection on male traits report moderate effect sizes (Jennions et al., 2012), with evidence that males are often under variable selection pressures within and across breeding seasons (Chaine & Lyon, 2008; Kingsolver et al., 2012; Robinson et al., 2008; Steele et al., 2011), have trait values near optima (Evans, 1998; Rodríguez et al., 2006), or that the magnitude of trait differences may not reflect the strength of current selection on males (Miller et al., 2016). Indeed, sex differences in signals can be caused by a range of selection pressures on both males and females, resulting in exaggeration or reduction of a variety of sex-specific signals (Bell & Zamudio, 2012; Dunn, Armenta, and Whittingham 2015; Price, 2015; Shultz & Burns, 2017; Wiens, 2001). To better understand how divergent selection operating between the sexes drives sex differences, we require holistic approaches that fully describe the traits of both males and females (Hare et al., 2019; Riebel et al., 2019). Ideally, studies should include multiple signaling traits that mediate inter- and intra-sexual interactions (Bro-Jørgensen, 2010; Hebets et al., 2016; Hebets & Papaj, 2004; Partan & Marler, 2005), as well as their fitness consequences.

Bird song and plumage color offer excellent examples for how female, as well as male, signal evolution can lead to sex differences. Females sang in the ancestor of modern songbirds and still sing in many tropical and temperate oscines (Garamszegi, et al., 2006; Odom et al., 2014; Price et al., 2009), suggesting that losses of female song have most often created sex-based differences in song. Similarly, in species in which females have dull plumage, there has been repeated, independent selection on feather color in both males and females (Dale et al., 2015; Hofmann et al., 2008; Price et al., 2014), with females experiencing more evolutionary plumage change than males in some lineages (Price et al., 2014). Even though losses of female song and elaborate plumage appear to contribute considerably to sex differences in signaling traits, the processes and conditions leading to these changes within individual species have not received much attention, particularly in species where female song is rare (Brunton et al., 2016; Kleindorfer et al., 2016).

Large-scale evolutionary change in dimorphic signaling traits is generally assumed to result because some signals increase fitness. Numerous studies have identified male plumage traits associated with strong female preferences and increased fitness (Byers et al., 2010; Møller, 1988; Rodríguez et al., 2006; Ryan et al., 2019; Searcy & Andersson, 1986). In contrast, relatively few studies have looked for or uncovered relationships between female visual signals and fitness proxies, but see: (Amundsen et al., 1997; Bulluck et al., 2017; Cain & Ketterson, 2011; Jawor et al., 2004; Pap et al., 2019). Likewise, many studies have examined the fitness correlates of male song (Catchpole & Slater, 2003; Gil & Gahr, 2002), but only a handful have looked at how female song impacts fitness (Brunton et al., 2016; Cain et al., 2015; Kleindorfer et al., 2016; Krieg & Getty, 2016). Although inclusive, systems-based approaches can best untangle the relative contributions of selection on males and females to signal evolution across modalities (Hebets et al., 2016; Riebel et al., 2019), researchers have seldom included both visual and acoustic phenotypes from both sexes within the same studies (Hebets et al., 2016; Riebel et al., 2019), but see (Webb et al. 2016). Moreover, to our knowledge, no study has directly tested whether the pattern of sex differences in vocal and visual sexual signaling traits is better predicted by contemporary selection on males (as is often assumed) or females. This likely reflects the difficulty of collecting reproductive success in the field, a lack of study in females, and a resulting paucity of datasets where the same traits and associated reproductive success are measured for both sexes. This is a particular challenge in species with infrequent female song, although such species provide a fertile testing ground for asking questions about why female song is reduced and differs from male song.

We propose that North American barn swallows (*Hirundo rustica erythrogaster*) offer a valuable system in which to investigate the selection pressures governing the evolution of sexual signaling traits and associated sex differences. We explored patterns of plumage and song in a population of male and female barn swallows by: 1) Describing female song in this species, 2) exploring the form and seasonal timing of song production in both sexes, 3) determining which acoustic and visual traits are robustly dimorphic, and 4) testing whether sex differences are more strongly associated with contemporary selection (i.e. trait correlations with reproductive success) in males or females. Because the differences between several measured male and female traits were not categorical, we conservatively discuss them as sex differences rather than dimorphism although other studies of this system may choose to call them dimorphic.

Much previous research is predicated on the idea that sex differences and dimorphism in communicative traits result from selection for male trait exaggeration (Badyaev & Hill, 2003; Freed, 2000; Langmore, 1998; Riebel, 2016; Riebel et al., 2005, 2019; Rosvall, 2011). Here, we test that assumption by including parallel data for both sexes and testing for linear associations between a measure of current selection and the magnitude of trait sexual differentiation in a variety of visual and acoustic traits (similar to Badyaev & Martin, 2000). If sex differences are driven/maintained by directional selection on males (for showier or more elaborate traits), we expect that trait correlations with male reproductive success will predict degree of sexual differentiation. However, if differences are driven/maintained by directional selection on females (for more cryptic or energy-efficient communication), we expect that trait associations with female reproductive success will predict levels of sexual differentiation. While the combined effect of differential selection on the sexes is ultimately responsible for the overall pattern of sex differences (Hedrick & Temeles, 1989; T. D. Price, 1984), we are chiefly concerned with testing the common assumption that it is sexual selection on males, rather than females, that drives this pattern in signaling traits. Alternatively, if trait sex differences arose in the past, but are currently under stabilizing selection, we would expect to find that the magnitude of trait sex differences have no contemporary associations with reproductive performance.

## METHODS

### Background

As the subject of hundreds of studies on sexual signal evolution since the late 1980s, barn swallows offer an excellent system for testing questions related to sex differences. Barn swallows are a weakly dimorphic species, in which males have darker ventral plumage and longer tail feathers on average. They comprise six Holarctic subspecies that have rapidly diversified from a common ancestor as recently as 7,700 years ago (Smith et al., 2018). In North American barn swallows (*H. r. erythrogaster),* males with darker ventral feathers have higher reproductive success, while tail feather length is not a preferred trait in males (Eikenaar et al., 2011; Safran & McGraw, 2004; Safran et al., 2016), or females (Safran & McGraw, 2004). There is limited evidence of selection for darker plumage in females in a New York population of *H. r. erythrogaster* (Safran & McGraw, 2004).

Previous barn swallow studies indicate that different components of male song (e.g. song rate and duration, rattle frequency and length) may provide receivers with information on the age, viability, condition, motivation and/or overall quality of singers (Dreiss et al., 2008; Galeotti et al., 1997, 2001; Garamszegi et al., 2005; Garamszegi, Hegyi, et al., 2006; Saino et al., 2003; Wilkins et al., 2015). Despite intensive study of this species, female signals have received relatively less attention. While there is a modest body of research into male song, there are no indexed papers describing female barn swallow vocalizations in any detail. Females have been reported to produce a “twitter-warble” song (Brown & Brown, 2020), but no formal descriptions or quantitative analyses of that song exist (see Supplementary Methods and Background for clarifying notes on female song).

### General Field Methods

Barn swallows in this study were part of a long-term study population in Boulder County, CO, which was set up in 2008 by Author 1 by contacting local birders and equestrian clubs, surveying the county for culverts and structures with old nests, and, after arrival, looking for signs of flying or perched swallows nearby. The five sites included in this study covered roughly 65 km^2^ and ranged from 8 to 64 banded individuals. Persistent mist netting effort throughout the breeding season resulted in the capture and unique marking of every or nearly every bird present at each study site (using color bands, and application of Sharpie marker ink combinations to left and right rectrices). These individual markings were used to identify individuals during sound recording. At the time of capture, tail streamer length was measured and a small sample (<90μL) of blood was taken via brachial venipuncture and stored in 2% lysis buffer for later parentage analysis. Additionally, a set of ~5 contour feathers were sampled along a ventral transect (throat, breast, belly, and vent), and attached to index cards with tape for color analysis using a spectrophotometer in the lab (Hubbard et al., 2017; Wilkins et al., 2015). Nests were closely monitored for egg-laying activity, with checks conducted at least every four days to determine clutch initiation date, and date of hatching. On approximately day 12 post-hatching, all nestlings in a nest were banded and blood samples taken for parentage analysis.

### Ethical Note

We made every possible attempt to minimize handling time and other sources of stress on swallows in our study population. Capture typically involved closing off all but one exit from a barn or other structure, in front of which a fine mesh nylon mist net was held or attached in place with bungee cords. Nets were constantly monitored and birds were usually removed within one minute of landing in the net and placed into custom-sewn cloth bags (closed with a ribbon) to calm them until they could be transported to a temporary banding station a short distance away (usually <30m), sampled, and released. Birds held temporarily in bags were kept in a quiet, cool location until they were removed for processing. Typically, birds are quiet and fairly still both in the net and while in the bag. Obvious signs of stress during handling were attended to constantly. In the rare instances where a bird showed signs of distress during netting, banding, measurement, or blood sampling, it was immediately released. All field methods were approved by the University of Colorado Institutional Animal Care and Use Committee (Protocols 07-07-SAF-01 and 1004.01).

### Visual Trait Measures

Consistent with a previous study on complex male signaling in this population (Wilkins et al. 2015), we measured maximum tail streamer length (rather than mean, as it is more normally distributed; See Table S1 for trait definitions). Additionally, we measured a total of 12 traits to characterize color variation: three axes of color (average brightness, hue, and red chroma) along a four-patch ventral transect (throat, breast, belly, and vent). We opted to use these huesaturation-brightness (HSB) measures to make interpretation and comparison to other studies in this system as easy as possible. We are also confident about the biological relevance of these trait measures, since they strongly correlate with eumelanin and phaeomelanin concentrations in feathers (McGraw et al., 2005), ventral feathers do not show an ultraviolet reflectance peak (Safran & McGraw, 2004), and color manipulations (measured using HSB) to mimic natural melanization profiles have shown predictable impacts on physiology and reproductive performance (Safran et al., 2005, 2008, 2016). However, as a check, we calculated color measures from raw spectra in the tetrahedral color space (TCS) that incorporates a model of avian visual sensitivity (Endler & Mielke, 2005; Stoddard & Prum, 2008). See Supplementary Methods for details on how we calculated both HSB and TCS color traits. As shown in Figure S2, average brightness is exactly equivalent to the TCS measure ‘brilliance’ described in Stoddard and Prum (2008). Our measures of chroma showed a strong positive (0.84) Spearman’s correlation with the equivalent TCS measure of saturation (rA, achieved chroma). Hue showed a moderate (−0.35) correlation with brilliance, though direct comparison with TCS measures of hue is challenging because TCS θ and Φ jointly describe stimulation of avian cones and a single correlation between hue and one of these measures in not meaningful. Overall, given biochemical and experimental evidence, and a lack of significant UV reflectance in barn swallow ventral coloration (Hubbard et al., 2017, McGraw et al., 2005), we are very confident that our HSB measures of color capture biologically meaningful variation in this system.

### Fitness Measures

We used female seasonal (i.e. annual) fledging success and male within-pair genetic paternity as fitness proxies for females and males, respectively. Female fledging success was calculated as the number of offspring leaving the nest across all broods. Genetic paternity was calculated as the number of fledged genetic offspring within the social nest across all broods, determined through genetic paternity exclusions. Paternity assignments (and total extra-pair paternity) were not feasible in our study area, due to the presence of known, unmonitored breeding sites within easy flying distance from monitored sites. To get a representative sample of males’ EPY would also require genotyping thousands of individuals in any given year. Briefly, genotypes were derived from fluorescent-labeled PCR products of six microsatellite loci. Paternity was conducted using CERVUS v3.03 (Kalinowski et al., 2007) and an offspring was considered extra-pair when the offspring-mother-father trio confidence did not reach the 95% level. Detailed paternity exclusion methods are in the Supplementary Methods. Four of the 12 fully-sampled males and females were social pairs, though we did not treat this statistically as the fitness correlational analysis was conducted separately for each sex.

### Recordings

Female barn swallows were recorded opportunistically as part of a study of male song (Wilkins et al., 2015) conducted in 2011 and 2012. Though males were the target of the prior study, this rarer, unpredictable female vocalization was of growing interest to Author 1. When a female vocalization was heard, Author 1 immediately redirected the microphone at the source of the sound, capturing as many vocalizations as possible. Given the complex acoustic environment in which barn swallows live, active, annotated recording is necessary for confident singer IDs. Thus, our samples by no means capture comprehensive singing outputs for individuals; however, our estimates of relative singing outputs should reflect realistic activity patterns. Songs were recorded in 16-bit WAV format, with 48kHz sampling rate using a Marantz PMD 660 paired with a Sennheiser MKH 20 and Telinga parabola (2011), or a Marantz PMD 661 paired with a Sennheiser ME62/k6 microphone and Telinga parabola (2012). Total recording time was approximately 57 hours (May 2-May 31) in 2011 and 48 hours (May 1-Aug 21) in 2012, all between 5am and noon (Wilkins et al., 2015).

### Acoustic Trait Measures

Using Raven v1.5 (Bioacoustics Research Program, 2011), we extracted acoustic parameters of all recorded songs from the 15 females (n=64) and a comparable number of songs (n=66) for 15 males (chosen randomly from among those with high quality song recordings) to create two acoustic datasets: (1) an element-level dataset of acoustic parameters measured for every element in all songs, and (2) a song-level dataset of acoustic parameters averaged for all elements in a song or calculated for the whole song. The element-level dataset was used to estimate an element diversity score (Keen et al., in review), which estimates the variability, or diversity, of elements for a given song within an acoustic feature space of all barn swallow element measurements (see full explanation in Supplementary Methods). This value was then appended to the song-level dataset, which was used for all subsequent statistical analyses involving song.

Of the 18 song-level variables extracted (see Supplementary Methods), we selected 11 that were likely to be robust to noise in recordings from barn swallow habitats and relevant for sexual signaling, based on previous studies of male song (Galeotti et al., 1997; Garamszegi, Hegyi, et al., 2006; Wilkins et al., 2015). The 11 selected acoustic variables were: song entropy (a measure of tonality), dominant frequency range, mean peak frequency, number of elements per song, element duration, element rate, song duration, gap duration, frequency interquartile range, and element diversity (see Table S1 for definitions). We then calculated individual means and coefficients of variation for each acoustic variable. This resulted in 22 variables describing differences in the magnitude and variability of different aspects of song frequency, timing, and elemental diversity for both sexes. The number of song traits was further reduced to avoid redundancy in analysis of sex differences and selection, as described below.

### Statistical Analysis

Our sample included 15 females with at least one song measured. To make female-male comparisons that are not skewed by uneven sample sizes, we randomly selected 15 males with high quality recordings from our data set. Our subsample of males was temporally equivalent to that for females, as the sexes did not differ in recording date relative to first clutch initiation: mean ± (SD) for females: −6.6 ± (10.7) and for males: −2.6 ± (21.1); t_unpaired_ = −0.65, df= 20.8, p= 0.52. For both sexes, due to missing data, we had 12 individuals with acoustic, visual, and fitness data. Because three females only had a single song recorded and coefficient of variation could not be calculated, only nine females had data for all 36 variables (22 acoustic traits,13 visual traits, and a fitness proxy). Thus, correlations between a particular trait and fitness metric in females were calculated from sample sizes ranging from nine to 15 individuals. Although our sample sizes are limited, the data set represents extensive population sampling time, and provides a starting point and methodology for more concerted effort.

All analyses were conducted using R v3.6.0 (R Core Team, 2019). To measure the magnitude of the sex difference for each trait, we used Cohen’s d, calculated as the difference of male from female population means divided by the pooled standard deviation (Cohen, 1988, p. 21). Thus, negative difference values indicate male-biased differences and positive values indicate female-biased differences. For example, in barn swallows, tail feathers show male-biased differentiation (as males have longer tails than females on average), resulting in a negative value on our sex difference scale. To determine which traits were robustly distinct, we created 10,000 bootstraps using the ‘bootstraps’ function in the {rsample} package v0.0.5 (Kuhn et al., 2019), resampling sets of 15 males and 15 females with replacement. We then calculated 95% confidence intervals from this posterior distribution using the base function ‘quantile.’

To avoid collinearity in further analysis of dimorphic traits, we identified variables with Spearman’s rank correlations >|0.7| and retained the most biologically intuitive variables for analysis. For example, average number of elements, song duration, and our measure of element diversity showed pairwise correlations >0.8. As element diversity is a higher level measure of complexity, for which there is a wide literature suggesting it to be a common target of selection (Benedict & Najar, 2019; Snyder & Creanza, 2019), we retained mean element diversity in our set of dimorphic traits, and discarded mean song duration and mean number of elements. After filtering out redundant and non-dimorphic traits, we were left with 10 visual and acoustic traits for exploring the connection between sex differences and contemporary selection pressures.

### Testing trait-wise associations between reproductive success and sexual differentiation

As a measure of contemporary selection pressures, we calculated each trait’s Spearman’s rank correlation with a fitness surrogate (seasonal number of fledged offspring for females and seasonal number of within-pair genetic offspring for males). Finally, in order to test the linear relationship between our surrogate measures of selection and observed trait sex differences, while accounting for nonindependence of traits, we used the ‘crunch’ function in the {caper} package v1.0.1 (Orme et al., 2018), which implements the CAIC (Comparative Analysis by Independent Contrasts) algorithm (Purvis & Rambaut, 1995). Following a similar approach to Garamszegi et al. (2006), we treated traits as species with known phylogenetic relationships (i.e. correlation structure). To build the tree used by ‘crunch,’ we calculated trait distances as 1 – abs(Spearman’s correlation), and constructed the tree using a single-linkage clustering method with the base ‘hclust’ function.

Thus, we made a linear model with a trait’s magnitude of sex difference as a response and a trait’s correlation with reproductive success as a predictor, controlling for correlation structure among traits. Given our metric of sexual differentiations, we would expect a significant positive association between a trait’s correlation with reproductive success and the magnitude of sexual differentiation if female fitness is maintaining sex-based trait differences. That is, traits that confer fitness benefits to females should have a high degree of differentiation for our metric, and traits that reduce fecundity in females should have low differentiation values. We would expect the opposite (a negative slope) if male fitness is driving/maintaining sexual differentiation. That is, traits associated with higher paternity would be more exaggerated in males (having more negative differentiation values on our metric), and traits that reduce paternity would be less exaggerated in males (higher differentiation values), resulting in a negative slope.

### Data availability

Male and female song clips, HSB and TCS color, and other phenotypic and fitness data are available in our GitHub repository at https://github.com/drwilkins/femaleSongInBARS.

Spectrograms in Figure 1 were generated with R code, available at https://github.com/drwilkins/rspect

**Figure 1.**
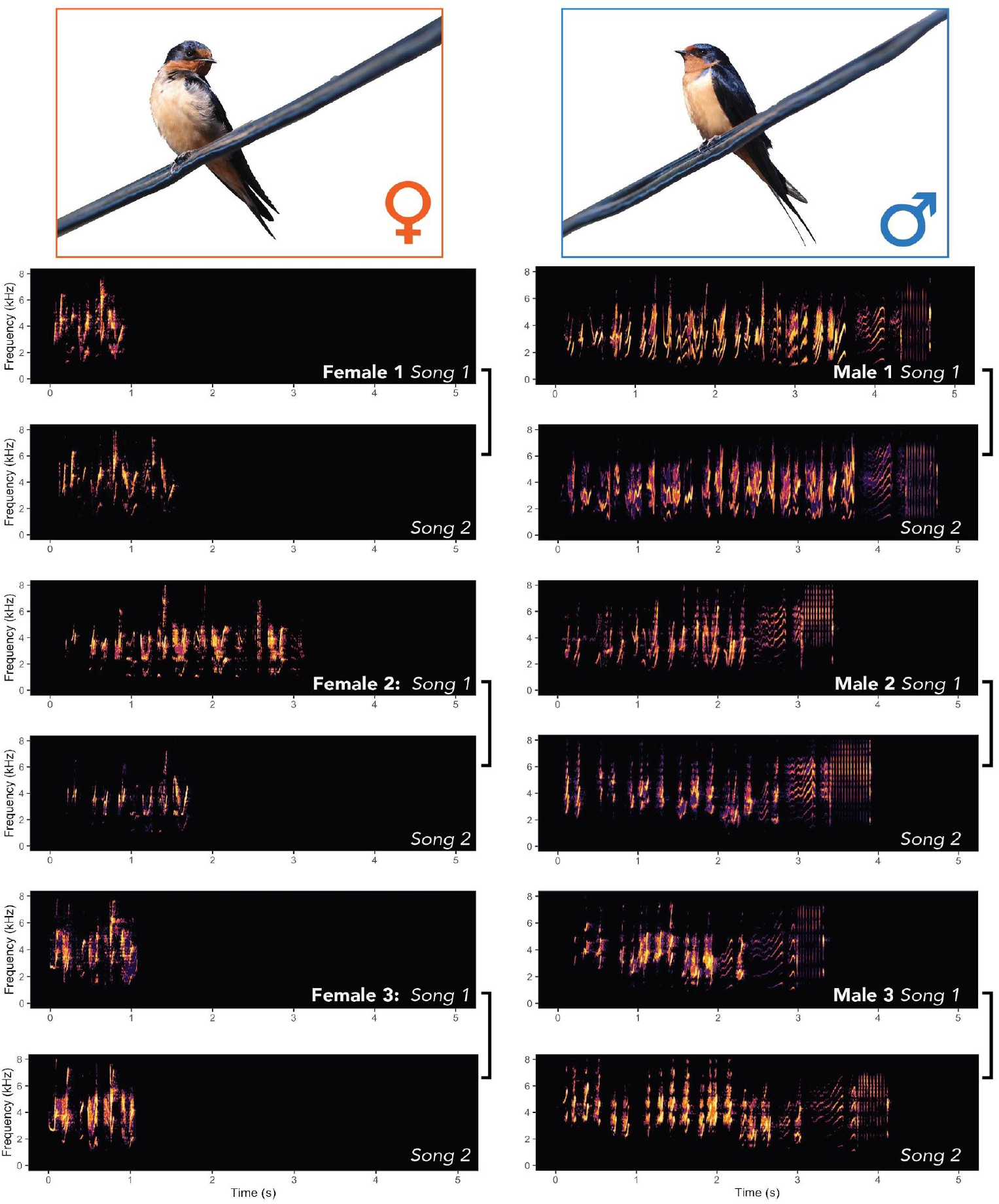
Spectrograms of female and male barn swallows. We show two song renditions for 3 females and 3 males to demonstrate variability. Photos by Author 1.

## RESULTS

### Characterizing female song

Recordings and observations revealed that females used a distinctive vocalization in spontaneous solo renditions, and in response to hearing other females, in a manner synonymous to male counter-singing. They were also observed using the vocalization to interrupt the songs of their mates (see supplementary video 1–vimeo.com/424642268). This vocalization is relatively complex, including many syllables with similar frequency modulation and pace to male song (Figure 1). Collectively, this evidence confirms that female barn swallows do produce facultative song, rather than just simple calls (Langmore, 1998).

Female song bouts were often produced within the nest, but also on perches near the nest or outside the barn/structure and were usually short and infrequent. As such, only 78 songs were recorded from 18 identifiable females over the course of two years, compared to 753 (865% more) songs from 40 males, given the same recording effort (~105 hours). This amounts to a recording efficiency of about 7.17 clear, identifiable songs/hr for males and 0.74 songs/hr for females over the course of the study. The sexes also differed strikingly in the phenology of song production. Male songs were observed and recorded between May 1st and August 21st, while female songs were only observed and recorded between May 10th and May 29th (pooling years) (Figure 2a). Restricting to only the active singing period for females during May of both years, a cumulative 91 hours of recording documented 8.9 individually identifiable songs/hr for males and 0.86 songs/hr for females. We could not calculate exact song rates per individual due to the complexities of recordings with shifting focal singers in dynamic colonial environments. To explore effects of breeding phenology on singing output, we also calculated relative recording dates by subtracting the ordinal date of first clutch initiation (i.e. breeding onset) from the ordinal recording date. For males, these relative recording dates ranged from 59 days before to 109 days after the first egg was laid by a males’ mate, with an average date (weighted by number of songs recorded each day) of 5.43 days *after* breeding onset; Figure 2b). For females the range was 29 days before to 14 days after breeding onset, with a weighted average of 3.90 days *before* breeding onset. Due to dropouts resulting from low recording quality, the total number of songs analyzed was 48 for females (mean=4.0, range= [1,11], sd=3.02) and 66 for males (mean=4.4, range= [2,5], sd= 0.99).

**Figure 2.**
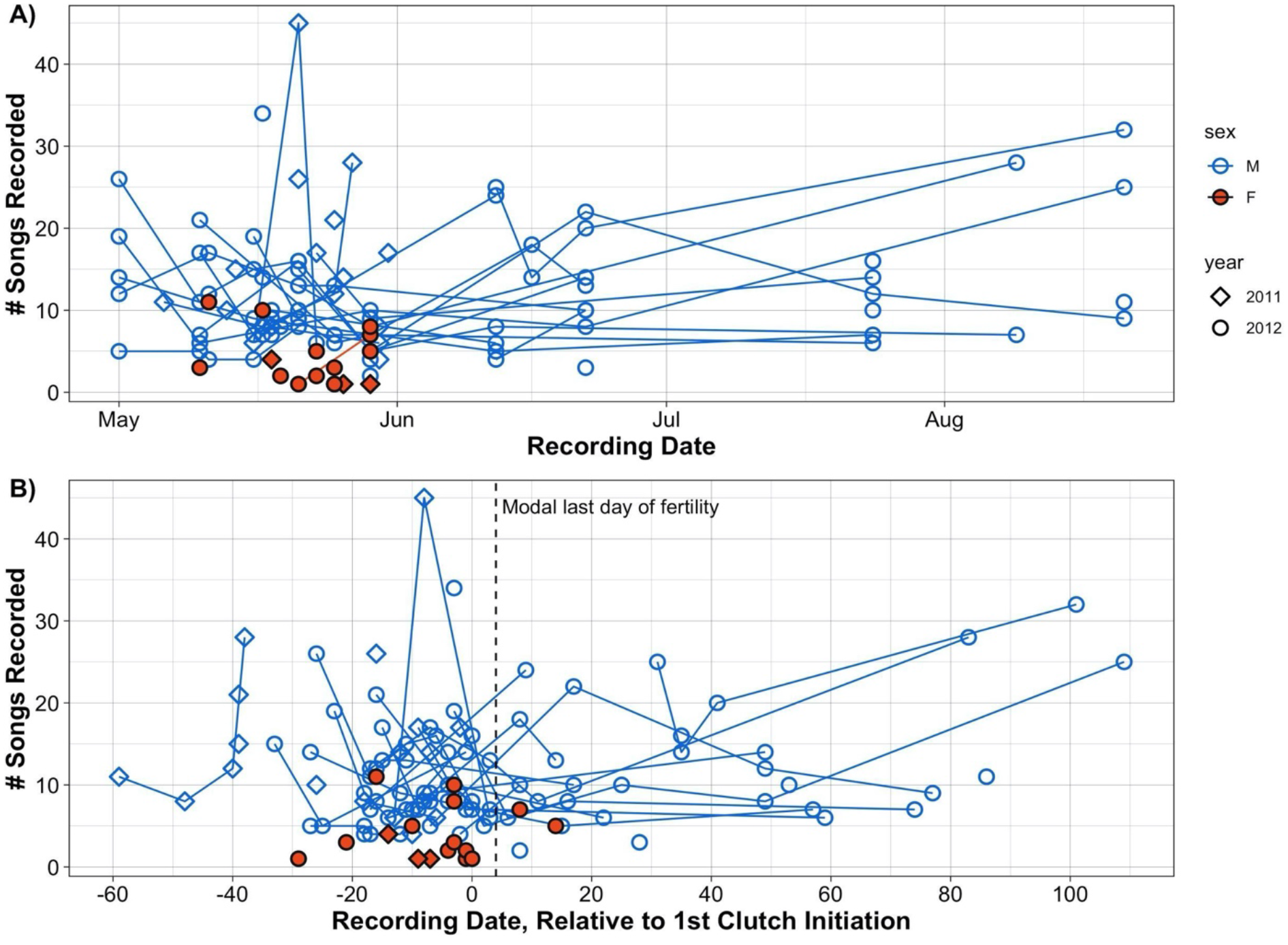
Seasonal variation in song production, as demonstrated by opportunistic recording of spontaneous male and female songs. A) Shows number of songs recorded for both sexes, as a function of recording date. In B) the ordinal date of breeding onset (first clutch initiation) has been subtracted from the ordinal recording date; thus, negative x-values represent recordings taken before breeding onset. The dashed vertical line marks Day 4 post clutch initiation–i.e. the last day of female fertility before laying of the ultimate egg of a modal 5-egg clutch. Lines connect points for individuals with recordings on multiple days.

### Visual and acoustic differentiation in males and females

Our bootstrap analysis of 35 acoustic and visual traits showed that 16 were robustly distinct in the two sexes-i.e. had bootstrap 95% confidence intervals that did not overlap zero (Figure 3). These include a mixture of song and visual traits, with song traits being most distinctive (13/22=59% for acoustic traits, compared to 3/13=23% for visual traits). Traits that had higher values in females (showing female-biased differences) included: entropy, CV_Mean Peak Frequency, CV_Dom Frequency Range, CV_Frequency IQR, CV_Element Diversity, Belly Brightness, and Element Duration. Male-biased dimorphic traits included: Vent Chroma, Frequency IQR, Mean Peak Frequency, CV_Entropy, Frequency Range, Tail Length, Element Diversity, Song Duration, and Element Count. Most notably, female songs are brief in duration, include few elements, include elements with high entropy (i.e. noise), and have low element diversity across songs, compared to males (Figure 3). Consistent with previous results for visual traits (Safran & McGraw, 2004), females had shorter tail streamers and lighter ventral plumage (higher brightness scores for belly and lower chroma scores for the vent). Trait means, sample sizes, and sex difference confidence intervals are reported in Table S2.

**Figure 3.**
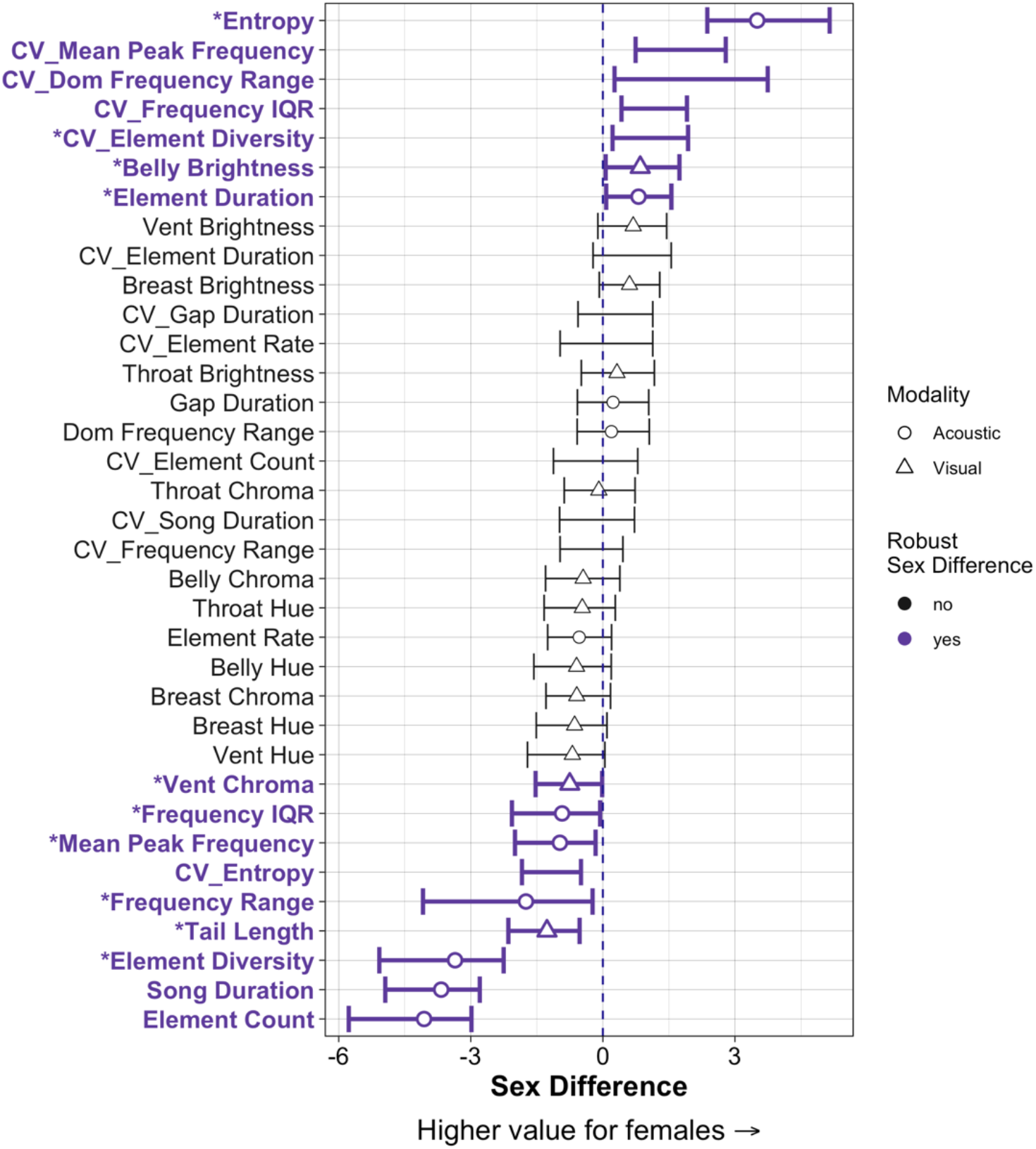
Sex differences in 35 potential sexual signaling traits. Empirical point estimates for Cohen’s d are shown, with 95% bootstrapped confidence intervals not overlapping zero bolded in purple. Symbols for points indicate trait modality (circle=acoustic, t triangle=visual). Traits are means, except where preceded by CV (coefficient of variation). Trait names beginning with a star were the chosen set of robustly sexually distinct traits used for further analysis, after removing highly redundant traits with Spearman’s correlations ≥|0.7|.

### Connecting contemporary selection to phenotypic sex differences

Our final objective was to test whether sexual differentiation is more strongly associated with contemporary selection in males or females (here, using surrogate measures of lifetime fitness: within-pair reproductive performance for a single breeding season). After controlling for intercorrelations between the 10 nonredundant, sexually distinct traits using independent contrasts, we found a significant positive relationship between trait sex differences and our measure of current selection pressures (i.e. correlation with either seasonal fledging success or genetic paternity) in females (df=8, slope=6.04, t=2.92, p=0.019), but not males (df=8, slope=1.06, t=0.547, p=0.599) (Figure 4). That is, traits that had a negative association with female reproductive success (see Figure S1) showed male-biased sex differences (i.e., greater trait expression in males), while traits that positively correlated with female reproductive success showed female-biased sex differences. The relationship remained significant for females when including all 16 robustly distinct traits (df=14, slope=4.93, t=3.53, p=0.003), as well as for the subset of distinct acoustic traits (df=11, slope=5.28, t=3.70, p=0.003). In contrast, the relationship between male traits and within-pair genetic paternity did not predict levels of sex differences for either of these subsets (all p≥0.260).

**Figure 4.**
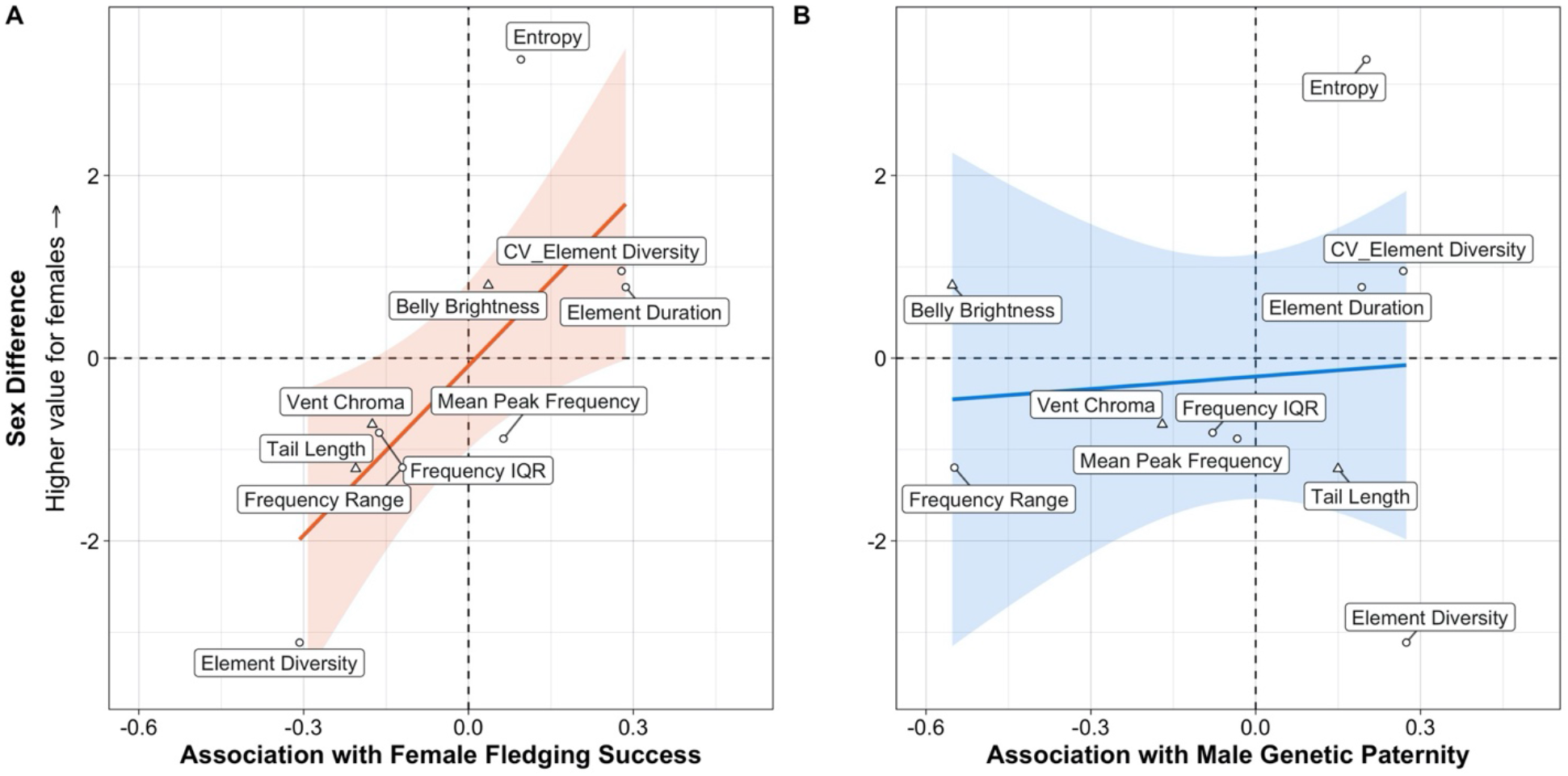
Relationship between trait sexual differentiation and trait association with reproductive output for A) females and B) males. Sexual differentiation was measured as the difference of scaled female and male trait values in pooled standard deviation units (Cohen’s d). Reproductive output was measured as seasonal fledging success for females and fledged within-pair seasonal genetic paternity for males. Females showed a much stronger (and significant) positive relationship between how traits impacted reproductive performance and the observed amount of sexual differentiation.

## DISCUSSION

### Female song in barn swallows

Despite intensive study of this species for decades, female song in barn swallows has been reported as absent by some authors (Sibley, 2014; Stokes & Stokes, 2010), the same as male song by others (Brown & Brown, 2020), or left ambiguous (Pieplow, 2017) (see Supplementary Methods for notes on previous descriptions). Our data and videos confirm that female barn swallows sing a quantitatively distinct song from males. We report that while female song is approximately ten times less frequent than male song and relegated largely to the first month of the breeding season, it has an overall similar structure to male song and seems to be used similarly in counter-singing, as well as mate-interruption contexts. The use of female song in counter-singing (see Supplementary Video 1) just prior to clutch initiation is consistent with singing patterns and use of song for territory/nest defense or intra-sexual competition in other Northern temperate breeding species (Cain et al., 2015; Krieg & Getty, 2016; Levin, 1996; Magoolagan et al., 2019; Rose et al., 2019; Yasukawa, 1989, 1990). Specifically, female barn swallow song could be actively maintained and used by females during this short period of the breeding season when females are establishing and competing for nest sites (Krieg & Getty, 2016; Rosvall, 2011). Further, the use of female song to interrupt male song fits with theories that describe song functionality in mate attraction, pair bonding, and mate signal jamming (Grafe et al., 2004; Tobias & Seddon, 2009).

Although female and male barn swallow songs have a generally similar form, female songs lack the terminal trill which is universal for male song among different barn swallow subspecies (Wilkins et al., 2018). Our results further indicate that female songs are shorter and noisier, with fewer, more variable elements, reduced frequency ranges, and lower element diversity (Figures 3, 4, S1). Thus, barn swallow song is categorically dimorphic in some aspects and shows variable sex differences in others. Overall, in this species song traits are more distinct by sex (59% of measured traits) than plumage traits (23% of measured traits) and the direction of sex differences is predicted by both the trait correlations with female fledging success and qualitative expectations from selection for efficient, cryptic signaling in females. For example, element diversity (our measure of average syllable complexity, which was retained after removing the redundant measure “song duration”) is much lower in females and showed a strong negative relationship with fledging success. In contrast, element duration (the length of individual song elements) was significantly longer in females and this trait showed the strongest positive correlation with reproductive success of any trait. Together, these results are consistent with selection for shorter songs with longer individual elements. Shorter songs may be favored to avoid attracting nest predators, an effect previously shown for song rate (Kleindorfer et al., 2016). Shorter songs also offer the cognitive benefit of simpler comparison due to easier discriminability of proportional differences (Akre et al., 2011; Akre & Johnsen, 2014).

The fact that mean element diversity is male-biased and negatively correlates with female fledging success, but CV_element diversity (i.e. song versatility: variability of syllable complexity across songs) is female-biased and positively correlates with fledging success was unexpected. One possible explanation for this may stem from variable functions (and audiences) for female song. That is, signals in competitive contexts tend to be shorter and more repetitive, which would select for low element diversity if this is the primary function of song (Collins et al., 2009; Galeotti et al., 1997). On the other hand, song versatility and the ability to adaptively change element diversity across competitive and mating-related signaling contexts could be especially important for females if they are constrained to shorter signal rate and duration, and there is a premium on efficient information transfer about motivation or quality within a shorter time window than males. This is, of course, speculative and close study of the signaling context of song production is necessary to better understand these patterns of female song variation.

Taken together, our results provide a mechanism supporting the evolution of female song in many North Temperate bird species from complex songs comparable to male songs toward highly condensed, context-dependent songs, or even toward the loss of songs altogether (Odom et al., 2014). This is significant because, while the vast literature on male birdsong (and acoustic signal evolution more broadly) has informed many aspects of sexual selection theory (Gil & Gahr, 2002; Nowicki et al., 2002), and male acoustic divergence has been shown to play a key role in premating isolation and speciation across diverse taxa (Alcaide et al., 2014; Blankers et al., 2019; Hasiniaina et al., 2020; Irwin et al., 2001; Lee et al., 2016; Sosa-López et al., 2016; Wilkins et al., 2013, 2018), we know relatively little about female vocal signaling or its implications for broader evolutionary processes, even in classic study systems like barn swallows.

### Overall patterns of sexual differentiation in signals

In contrast to a common assumption in the sexual selection literature, we found that overall patterns of sex differences are better explained by (surrogate measures of) selection on females, rather than selection for more elaborate males. That is, the overall magnitudes of sex-based differences in song form, tail streamer length, and color were predicted by how those traits correlated with female, but not male reproductive performance. This does not mean that selection on males (i.e. through female choice) is not important or relevant in explaining sex differences. For example, as shown in Figures 4B and S1, belly brightness-a known sexual signal in males within this population (Safran et al., 2016; Wilkins et al., 2015)–showed a strong negative association with genetic paternity (selecting for darker males with lower brightness). In turn, belly brightness in females showed a slight positive association with fledging success, and sex difference was low (0.8 SD), but significantly biased towards lighter females. That is, higher reproductive performance among darker males, combined with negligibly higher performance in lighter females correlates with an overall pattern of slightly darker males. In contrast, a strong positive correlation between element diversity and reproductive success in males and a strong negative correlation for females (i.e. divergent selection in the sexes) corresponds with a highly male-biased sex difference (−3.11 SD) toward more complex syllable composition of songs. While an additional analysis testing the combined impact of trait-wise reproductive performance on trait-wise sex differences for both sexes is desirable, it would likely be similar to the result shown for females (Figure 4A) and is not possible using our current comparative analysis by independent contrasts approach, as we are unaware of a way to control for known differences in trait correlation structure between the sexes in a single model.

### Broader implications

Although our small sample size limits our ability to rigorously estimate selection gradients or account for error in our estimate of the correlation between traits and fitness proxies, our findings offer baseline evidence that selection on females may be more consistent and therefore more important in maintaining phenotypic distinctions between the sexes within populations. This is perhaps unsurprising, given previous studies showing variability in female preferences for male traits (Chaine & Lyon, 2008; Kingsolver et al., 2012; Robinson et al., 2008; Steele et al., 2011). Nevertheless, it has important implications for comparative studies which rely on the degree of plumage differences as a surrogate for the strength of sexual selection. It is widely recognized that there are limitations to using sex differences, including dimorphism, as a surrogate for sexual selection (Bell & Zamudio, 2012; Huang & Rabosky, 2014; Kraaijeveld et al., 2011; Price, 2015). We would add the caveat that researchers must consider the action of sexual selection on both males and females, and include the traits of both sexes, even if traits are reduced or difficult to sample. Many studies assume that sex differences largely result from sexual selection for exaggerated male traits in conjunction with ecological selection for crypsis on females. This may be a valid assumption in some cases, such as in sexual dichromatism in damselflies (Svensson & Waller, 2013). However, our results highlight the need (especially in taxa with more complex communication and/or cognition) to consider how female competition and mutual mate choice affect the evolution of sex differences within the context of constraining ecological selection on females from increased costs of predation, migration, and/or coloniality (Jawor et al., 2004; Price, 2015; Tobias et al., 2011). Outcomes of this study suggest that following initial evolution of a mutually ornamented ancestor, sex differences and dimorphism in a broad suite of visual and acoustic traits was most likely created and maintained in barn swallows through countervailing selection on females, rather than on males (Kraaijeveld et al., 2007).

## CONCLUSIONS

With accumulating evidence for mutual mate choice and mutual ornamentation as the ancestral state across diverse taxa (Edward & Chapman, 2011; Kraaijeveld et al., 2007; Odom et al., 2014; Hofmann et al., 2008), we offer a method for testing whether selection on females may promote sex differences to a much greater extent than is currently appreciated. Collectively, our results highlight the importance of studying rare behavioral phenomena and, in particular, call for better documentation and targeted research on female song (Karan J. Odom & Benedict, 2018). There are growing accounts of temperate-zone female birds that sing for short time periods during the breeding season, particularly during settlement of breeding territories (Hathcock & Benedict, 2018; Krieg & Getty, 2016; Taff et al., 2012). Thus, more focused research effort during this time period may very well show that complete loss of song in female Passerines is the exception, rather than the widely purported rule, and that similar selective pressures shape female song in multiple species.

Though there is increasing evidence that female signaling traits can evolve as fast or faster than male traits across species (Johnson et al., 2013; Dunn, Armenta, and Whittingham 2015; J. J. Price, 2019), we have few studies within species to guide our understanding of the processes driving these patterns. In contrast to the vast literature on within-species geographic divergence in male signals (McLean & Stuart-Fox, 2014; Podos & Warren, 2007; Slabbekoorn & Smith, 2002; Velásquez, 2014), there are few studies for females in the visual modality (McCoy et al., 1997; McLean & Stuart-Fox, 2014; Obara et al., 2008; Roulin, 2003; Tuomaala et al., 2012) and fewer still in the acoustic modality (Graham et al., 2017, 2018; Mennill & Rogers, 2006; Odom & Mennill, 2012). Yet, it is worth noting that three studies (in butterflies and birds) that considered intraspecific geographic variation in both sexes found greater signal divergence in females than males across populations (Graham et al., 2018; Mennill & Rogers, 2006; Tuomaala et al., 2012). The implications of female signal evolution, male mate choice/recognition, and their impact on speciation have received little attention in the literature, with a few relevant studies focusing primarily on fish and insects (Chung et al., 2014; Edward & Chapman, 2011; Jiggins et al., 2004; Roberts & Mendelson, 2017, 2020). Targeted study of both sexes in concert across taxa is necessary to gain a more holistic understanding of how signals evolve within and among populations and how these trait changes feed into larger ecological, evolutionary, and eco-evolutionary processes (Bonduriansky, 2011; Cole & Endler, 2015; Endler & Basolo, 1998; Fryxell et al., 2019).

## ACKNOWLEDGEMENTS

MRW thanks the Macaulay Library for loaning him audio equipment in 2011, and numerous undergraduate and graduate students in the Safran Lab who helped collect the mountain of phenotypic and fitness data that enabled this study. He would also like to thank the many site owners who allowed us to traipse around their property at odd hours of the day (in particular, the Colorado Horse Rescue). We also thank Megan Barkdull for help with sound analysis and Joey Hubbard for sage counsel on color analysis.

## FUNDING

This work was supported by an American Ornithological Society: Joseph Grinnell Award and an NSF Graduate Research Fellowship to MRW. KJO was supported by a U.S. National Science Foundation Postdoctoral Research Fellowship in Biology (grant no. 1612861).

## Electronic Supplement

**Video S1**. “Female Barn Swallows Sing!”: provides detail and examples about timing and potential functions of female song. Accessible at: https://vimeo.com/424642268

### Supplementary Methods and Background

#### Acoustics

##### Clarifying notes on female song

Popular field guide entries for barn swallows do not specify which sex sings, and it can be assumed the song description refers to males (Sibley, 2014; Stokes & Stokes, 2010). In contrast, the *Peterson Field Guide to Bird Sounds of Eastern North America* states “females also reported to sing” (Pieplow, 2017) and the entry in *Birds of the World* explicitly states that both sexes sing the “twitter-warble” song, i.e. typical male song (Brown & Brown, 2020). These latter interpretations may stem from Samuel’s description of the barn swallow vocal repertoire, in which he states that the courtship song “is occasionally given by the female” (Samuel, 1971). He also describes a “whine call,” which may be the same as the copulation call (“waeae-waeae”) referred to in the *Birds of the World* (Brown & Brown, 2020), which may refer to what we call song in this study. In lieu of the original recordings or a high-quality spectrogram, it is difficult to assess which female vocalizations are being described in previous studies. In Møller’s *Sexual Selection and the Barn Swallow* (1994) there is no mention of female song or vocalizations at all. In fact the copulation call is ascribed to the male (p78). We believe this vocalization, the main phrase of which we transliterate as “dwidle dwight,” is more accurately described as a song rather than a call (sensu Marler, 2004), since it is multisyllabic, structurally complex, and seems to have a function in mate attraction, pair bonding, and competition, based on observations (see supplementary video).

##### General acoustic analysis details

Prior to song analysis, all songs were standardized to a sample rate of 44.1 kHz and a bit depth of 16. We used Raven 1.5 (Bioacoustics Research Program, 2011) to select and extract measurements for each element in each song. We defined an element as a single, continuous trace on a spectrogram, separated from other elements by a visible break in time or an extreme shift in overall harmonic structure (occasionally, structurally distinct barn swallow elements were separated by very short breaks in time, so this additional criterion helped us distinguish among elements). For pure tone elements, we selected and compared the fundamental frequency of the element, whereas for harmonically rich elements, we selected the fundamental to the dominant frequencies. We did substantial visual checking to be sure that elements were separated and measured consistently across all songs. We used the following Hann window settings with a sample size of 512 (DFT), hop size of 256, and 50 percent overlap in Raven v1.5 for selecting and measuring elements (Bioacoustics Research Program, 2011).

##### Element-level acoustic parameters

For every element in all songs we extracted a large number of energy-based measurements calculated from the cumulative amplitude of the spectrum or envelope using Raven Bioacoustic Analysis Software v1.5 and the warbleR package in R (Araya-Salas & Smith-Vidaurre, 2017; Bioacoustics Research Program, 2011). We removed any highly correlated variables with a Pearson correlation coefficient greater than 0.95. This resulted in the following acoustic variables measured for each element using each bioacoustics program: Raven measurements – peak frequency, frequency 5%, delta frequency, interquartile range bandwidth, time 5%, interquartile range duration, duration 90%, aggregate entropy, average entropy, average power, energy; warbleR measurements – standard deviation of frequency, mean dominant frequency, mean peak frequency, minimum dominant frequency, maximum dominant frequency, 1st quartile frequency, interquartile frequency range, dominant frequency range, dominant frequency slope, 1st quartile time, spectral entropy, time entropy, entropy, spectral flatness, modulation index, skewness, and kurtosis. See Raven and warbleR manuals for definitions of each parameter (Araya-Salas & Smith-Vidaurre, 2017; Bioacoustics Research Program, 2011).

###### Song-level acoustic parameters

Using the warbleR package in R, we extracted the following acoustic parameters for each song: mean frequency, mean peak frequency, median frequency, frequency range, dominant frequency range, 1st quartile frequency, 3rd quartile frequency, interquartile frequency range, median time, 1st quartile time, 3rd quartile time, entropy, average element duration per song, song duration, number of elements, element rate, and gap duration. See the warbleR vignette for descriptions of each parameter (Araya-Salas & Smith-Vidaurre, 2017).

###### Element diversity calculations

Using the element-level acoustic parameters, we estimated element diversity for each song as the area encompassing the elements for that song in a multi-dimensional feature space. To do this, we first created an element-level acoustic space by inputting the element-level acoustic parameters for every element for all songs into an unsupervised random forest. We used the ‘randomForest’ function in the R package {randomForest} with the following specifications: 10000 trees, minimal node size of 1, Gini impurity index as split rule, 5 randomly sampled variables at each split and out-of-bag proximity (Liaw & Wiener, 2002). This produced a proximity matrix, which we transformed into a set of five vectors using classic multidimensional scaling, using the ‘cmdscale’ function in the {stats} R package. We then calculated and extracted minimum convex polygons to calculate the area surrounding all elements for each song within this multidimensional space using the function ‘mcp’ in the {adehabitatHR} R package (Calenge, 2006). Therefore, each area is an estimate of the variability, or diversity, of elements for a given song within an acoustic feature space of all barn swallow element measurements.

#### Color Analysis

##### Hue Saturation Brightness (HSB) Color Measures

###### Methods reprinted from Wilkins et al. (2015), Online Appendix S1

Feather samples from four ventral patches (throat, breast, belly, vent) were taped to a standard white card background so that they overlapped as they do on the body of a bird. The color of each patch was measured using a spectrometer (USB 4000, Ocean Optics), pulsed xenon light (PX-2, Ocean Optics) and SpectraSuite software (v2.0.151). The probe was held at 90 degrees to the feather surface at a distance such that a 2.5 mm diameter of the surface was illuminated and measured. Each sample was measured three times, lifting the probe between measurements, and averaged. Each measurement was an average of 20 scans of the spectrometer. From the generated spectra, we calculated standard color descriptors: i) average brightness, ii) hue, and iii) red chroma. Average brightness was calculated as the average percent reflectance between 300 and 700 nm, hue was calculated as the wavelength that corresponds to where the slope of the curve is steepest between 550 and 700 nm, and red chroma was calculated as the proportion of light reflected in the red range (600 to 700 nm) relative to the entire range (300 to 700 nm).

###### Comparison of HSB to TCS color metrics

Though we have opted to use HSB methods for the current manuscript for easier comparison to most published barn swallow studies, we also calculated color measures in the tetrahedral color space (TCS) for comparison. TCS measurements follow Wilkins, et al. (2016). Briefly, we used the ‘tcs’ function in the R package {pavo} v. 0.5-5 (Maia et al. 2013). These color metrics describe the relative stimulation of the four bird cone channels (u, s, m, and l), relative to the achromatic origin. Thus, each color can be described by a vector and represented by θ (horizontal angle), Φ (vertical angle), and r (vector length), with θ and Φ reflecting hue. Because the color space is a tetrahedron, rather than a sphere, the maximum value of r varies by hue. As such, we use rA (achieved chroma, r/rmax) as a relative measure of saturation. Additionally, we calculated brilliance, or the averaged reflectance values for each measured wavelength between 300 and 700 nm. Collectively, our color metrics, θ, Φ, rA, and brilliance describe the color of each ventral plumage patch, quantified using the average avian UV visual model, as defined in {pavo}.

To assess the match between traditional HSB color metrics and the avian visual model color measures, we explored Spearman’s rho correlations between all color metrics calculated for the breast patch for the 27 individuals in our study with complete feather samples. TCS brilliance is synonymous with HSB average brightness. HSB chroma is very strongly correlated with the most relevant TCS metric (rA: achieved chroma). Hue shows a moderate negative correlation with TCS brilliance, though a direct comparison to TCS hue is complicated, due to two angular measures (θ and Φ) of hue in TCS space. Overall, the HSB and TCS show strong correspondence, suggesting that our reported measures of color represent discernible color variation for live birds, which is supported by experimental and biochemical evidence.

#### Paternity Analyses

##### Methods reprinted from Wilkins et al. (2015), Online Appendix S1

The six microsatellite markers used for paternity analyses were Escu6: (Hanotte et al. 1994); Ltr6: (McDonald & Potts 1994); Pocc6: (Bensch et al. 1997); and Hir11, Hir19, and Hir20: (Tsyusko et al. 2007)). Reaction conditions for pooled Escu6, Ltr6, Hir20, and Hir11 primers consisted of a 10 ul solution with 50-100 ng DNA, 0.12 mM of each labeled forward primer, 0.12 mM of each reverse primer, 200 M each dNTP, 3.25 mM MgCl_2_, 1x PCR Buffer, 0.15 units Taq polymerase (New England Biolabs, Massachusetts, U.S.A.), and were amplified with the following protocol: initial denaturation step of 94°C for 1 minute, followed by 10 cycles of 94°C for 30 s, 55°C for 30 s, and 72°C for 45 s, with an additional 25 cycles starting at 87°C for 30 s instead of 94°C, and completed with a final extension at 72 °C for 3 min. The Pocc6 reaction was modified from the above conditions by using 1.25 mM MgCl_2_, and modified for the Hir19 reaction with 3 mM MgCl_2_ and 0.2 mM of each forward and reverse primer. The PCR amplification protocol for Pocc6 and Hir19 was similar to the pooled loci protocol with the exception that 60°C was used for the annealing temperature. Amplified PCR products containing the fluorescently-labeled forward primer were detected using an ABI3730 DNA analyzer (ABI, Inc.). Allele peaks were manually called using Peak Scanner v1.0 (Applied Biosystems) in order to minimize genotyping error associated with irregular PCR products. For the paternity analysis simulation in CERVUS, we left the proportion of loci typed at 0.959 (determined from our data) and the proportion of loci mistyped at the default 0.01, and ran the program for 10,000 iterations. For each male, we set the female seen brooding eggs as the known mother and considered an offspring as extra-pair when offspring-mother-father trio confidence did not reach the 95% level. We adopted this approach, rather than a threshold number of parent-offspring mismatches for paternity exclusions because confidence levels are derived from likelihood equations which take into account potential genotyping errors.

---

**Figure S1.**
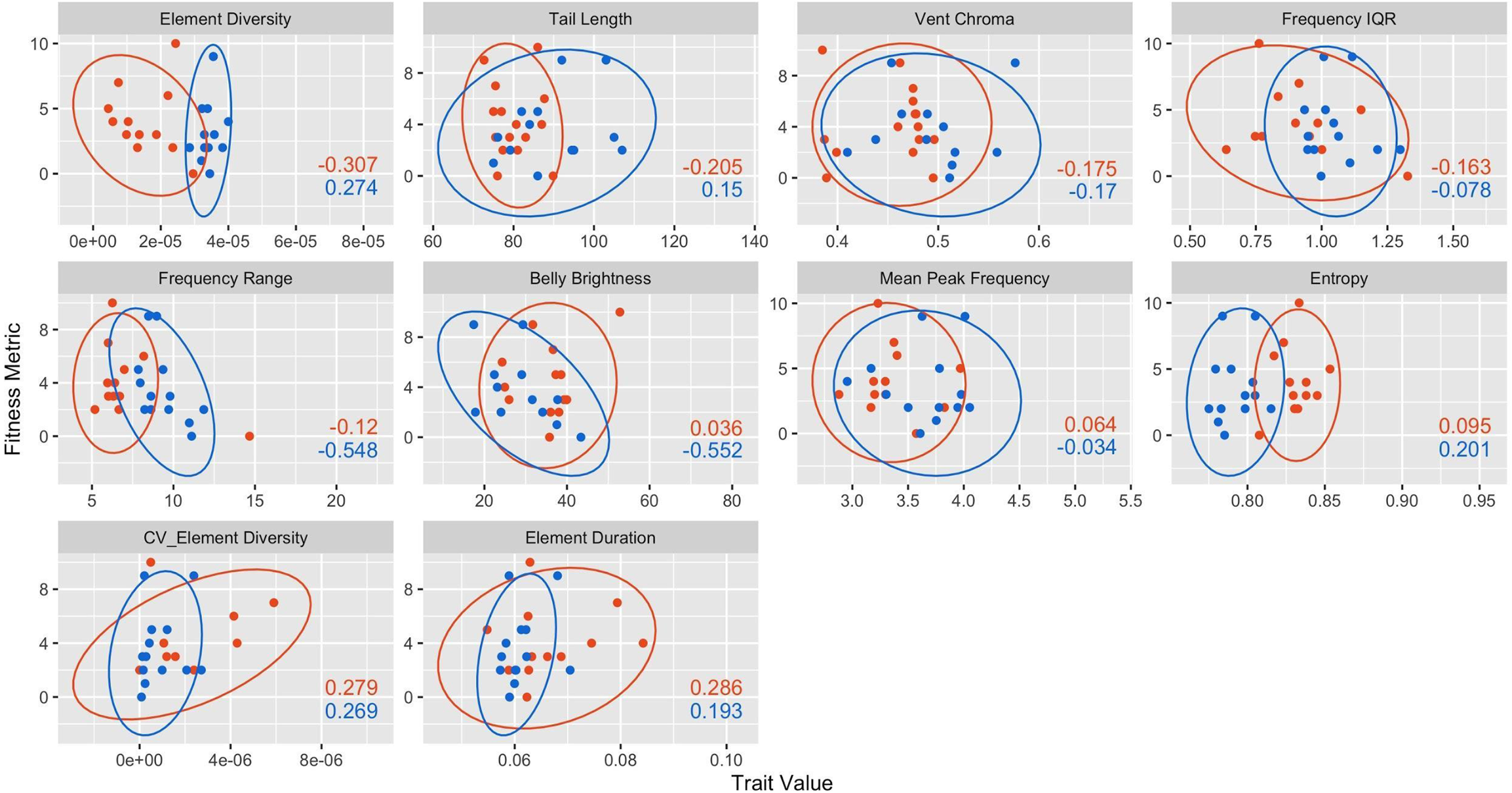
Spearman’s rank correlations between robustly dimorphic traits shown in Figure 4 and a relevant fitness metric–our measure of contemporary selection. The fitness metric for males was seasonal within pair genetic paternity, and for females it was seasonal fledging success. Orange 95% confidence ellipses and points are for females; blue ellipses and points are for males. Plots are ordered (top left to bottom right) from lowest to highest correlation for females.

**Figure S2.**
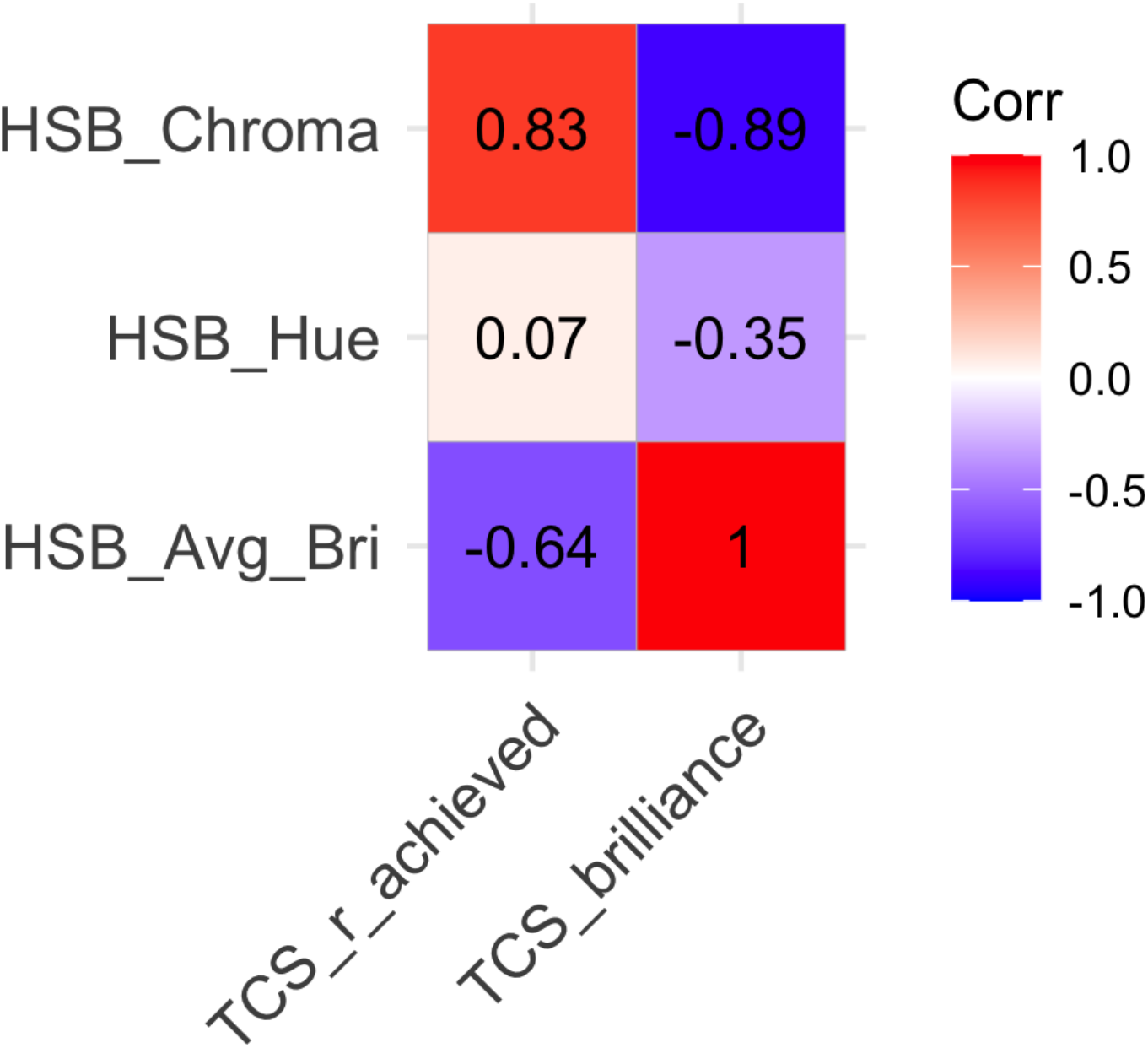
Spearman’s correlations between our reported Hue-Saturation-Brightness (HSB) color measures and tetrahedral color space (TCS) color metrics. All measures are for the 27 individuals (both sexes) in our dataset with complete feather color measures. Only the breast patch is shown, since correlations

**Table S1.**
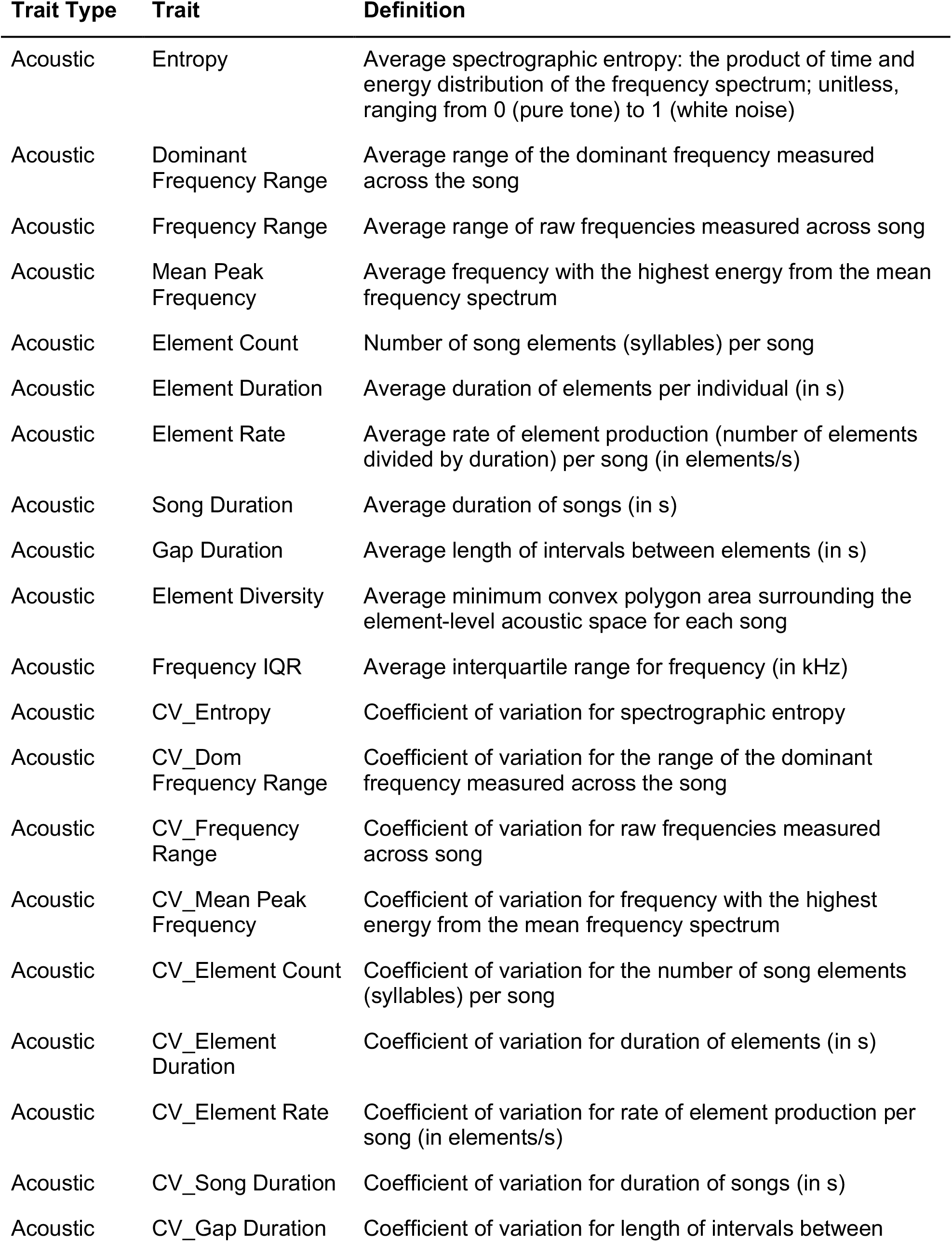
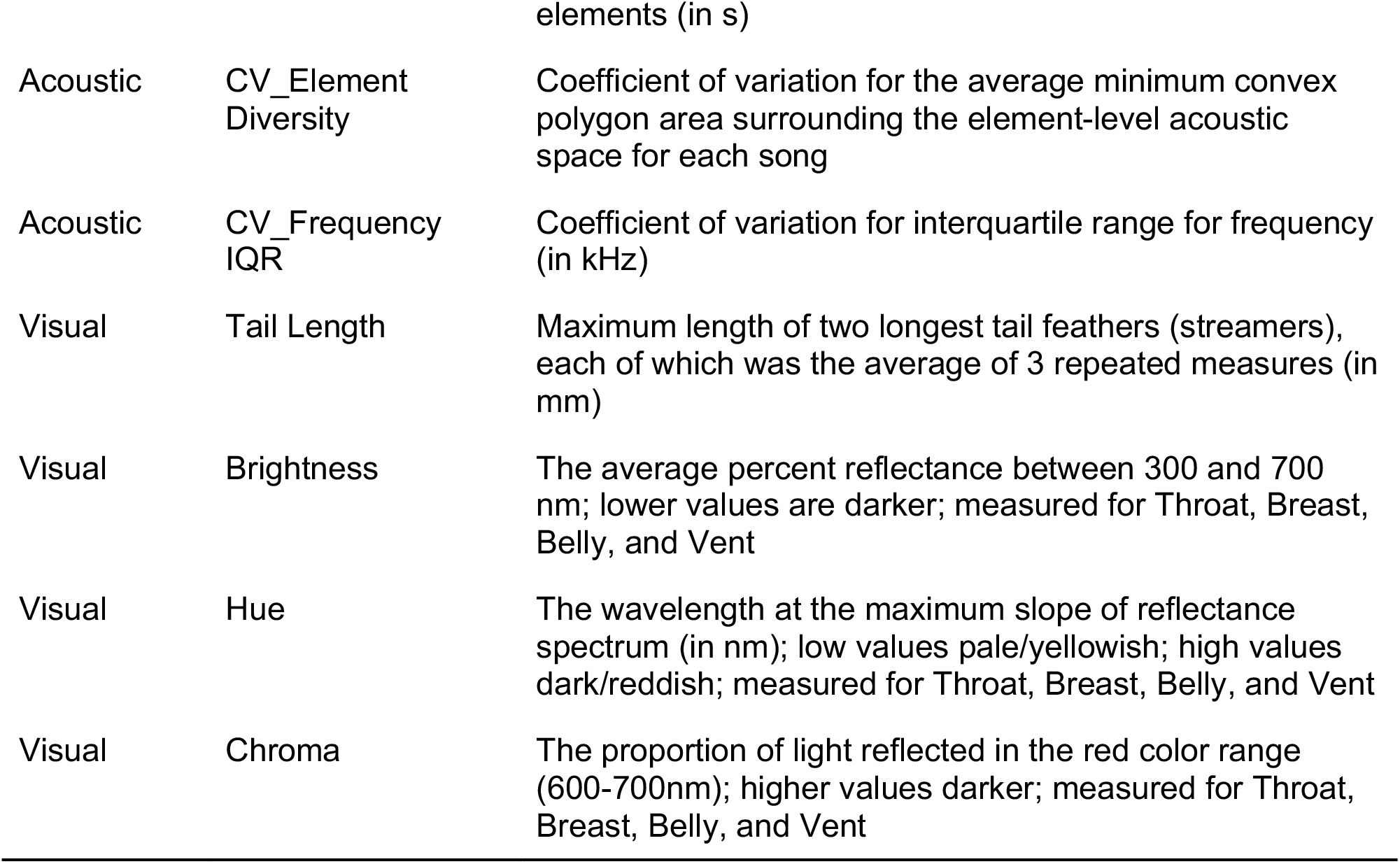
Trait definitions for all per individual visual and acoustic variables calculated for this study.

**Table S2.** Female and male, sample sizes, means, standard deviations, sex difference (i.e. Cohen’s D), and dimorphism 95% confidence intervals for 10,000 bootstraps. Sample sizes for coefficients of variation are lower for females, as three individuals only had a single song recorded. Bolded traits had dimorphism confidence intervals that did not overlap zero. Starred traits were selected for the final analysis in Figure 4, after eliminating the less biologically intuitive trait for redundant trait pairs with >|0.7| correlations.

| Trait | F_n | F_Mean (SD) | M_n | M_mean (SD) | Dimorp<br>hism | 95% Conf Int |
| --- | --- | --- | --- | --- | --- | --- |
| **\*Belly Brightness** | **13** | **35.5 (7.62)** | **14** | **29.3 (7.62)** | **0.799** | **(0.0694, 1.31)** |
| Belly Chroma | 13 | 0.421 (0.0434) | 14 | 0.441 (0.0434) | -0.425 | (-1.09, 0.333) |
| Belly Hue | 13 | 608 (29) | 14 | 626 (29) | -0.554 | (-1.21, 0.19) |
| Breast Brightness | 15 | 35.7 (9.39) | 14 | 30.8 (9.39) | 0.589 | (-0.114, 1.1) |
| Breast Chroma | 15 | 0.424 (0.0356) | 14 | 0.446 (0.0356) | -0.577 | (-1.1, 0.129) |
| Breast Hue | 15 | 616 (23.3) | 14 | 631 (23.3) | -0.62 | (-1.16, 0.0884) |
| **CV_Dom Frequency Range** | **9** | **0.0902 (0.0483)** | **15** | **0.0363 (0.0483)** | **1.27** | **(0.362, 1.79)** |
| CV_Element Count | 9 | 1.5 (2.24) | 15 | 1.41 (2.24) | 0.0468 | (-0.962, 0.875) |
| **\*CV_Element Diversity** | **9** | **2.34e-06 (2.01e-06)** | **15** | **8.61e-07 (2.01e-06)** | **0.954** | **(0.181, 1.57)** |
| CV_Element Duration | 9 | 0.000904 (0.000889) | 15 | 0.000465 (0.000889) | 0.626 | (-0.264, 1.34) |
| CV_Element Rate | 9 | 0.149 (0.188) | 15 | 0.0899 (0.188) | 0.432 | (-0.903, 1.15) |
| **CV_Entropy** | **9** | **0.000127 (0.000117)** | **15** | **0.000359 (0.000117)** | **-1.02** | **(-1.29, -0.454)** |
| **CV_Frequency IQR** | **9** | **0.069 (0.066)** | **15** | **0.0189 (0.066)** | **1.04** | **(0.517, 1.58)** |
| CV_Frequency Range | 9 | 0.25 (0.573) | 15 | 0.497 (0.573) | -0.319 | (-0.82, 0.582) |
| CV_Gap Duration | 9 | 0.00152 (0.0015) | 15 | 0.000986 (0.0015) | 0.454 | (-0.546, 1.15) |
| **CV_Mean Peak Frequency** | **9** | **0.0499 (0.0408)** | **15** | **0.0099 (0.0408)** | **1.37** | **(0.801, 1.77)** |
| CV_Song Duration | 9 | 0.117 (0.139) | 15 | 0.151 (0.139) | -0.199 | (-0.829, 0.774) |
| Dom Frequency Range | 12 | 2.21 (0.303) | 15 | 2.16 (0.303) | 0.172 | (-0.608, 0.958) |
| **Element Count** | **12** | **15.4 (6.28)** | **15** | **43.6 (6.28)** | **-3.79** | **(-1.88, -1.65)** |
| **\*Element Diversity** | **12** | **1.53e-05 (8.21e-06)** | **15** | **3.44e-05 (8.21e-06)** | **-3.11** | **(-1.88, -1.53)** |
| **\*Element Duration** | **12** | **0.0667 (0.00862)** | **15** | **0.0615 (0.00862)** | **0.778** | **(0.0664, 1.28)** |
| Element Rate | 12 | 10.3 (0.914) | 15 | 10.7 (0.914) | -0.516 | (-1.16, 0.27) |
| **\*Entropy** | **12** | **0.831 (0.0121)** | **15** | **0.792 (0.0121)** | **3.27** | **(1.54, 1.85)** |
| **\*Frequency IQR** | **12** | **0.916 (0.189)** | **15** | **1.04 (0.189)** | **-0.816** | **(-1.45, -0.0324)** |
| **\*Frequency Range** | **12** | **7.11 (2.49)** | **15** | **9.46 (2.49)** | **-1.2** | **(-1.77, -0.264)** |
| Gap Duration | 12 | 0.0347 (0.00477) | 15 | 0.0338 (0.00477) | 0.21 | (-0.583, 0.994) |
| **\*Mean Peak**<br>**Frequency** | **12** | **3.37 (0.299)** | **15** | **3.64 (0.299)** | **-0.881** | **(-1.45, -0.125)** |
| **Song Duration** | **12** | **1.49 (0.543)** | **15** | **4.12 (0.543)** | **-3.45** | **(-1.85, -1.62)** |
| **\*Tail Length** | **15** | **80.2 (5.36)** | **15** | **90.5 (5.36)** | **-1.21** | **(-1.45, -0.563)** |
| Throat Brightness | 13 | 22.2 (5.81) | 14 | 20.5 (5.81) | 0.3 | (-0.438, 1) |
| Throat Chroma | 13 | 0.482 (0.0443) | 14 | 0.486 (0.0443) | -0.0779 | (-0.825, 0.644) |
| Throat Hue | 13 | 641 (33.9) | 14 | 654 (33.9) | -0.458 | (-1.09, 0.327) |
| Vent Brightness | 15 | 27.7 (7.2) | 14 | 23.5 (7.2) | 0.651 | (-0.0368, 1.19) |
| **\*Vent Chroma** | **15** | **0.454 (0.0413)** | **14** | **0.487 (0.0413)** | **-0.725** | **(-1.21, -0.0361)** |
| Vent Hue | 15 | 628 (23.4) | 14 | 645 (23.4) | -0.642 | (-1.28, 0.0747) |

## Notes

https://github.com/drwilkins/femaleSongInBARS

https://vimeo.com/424642268

### Competing Interest Statement

The authors have declared no competing interest.

### Summary of Updates

Title and a decent amount of the text have been changed following final revisions before acceptance at Animal Behaviour. Specifically, the associate editor requested we change many usages of "dimorphism" to "sex differences".

